# INgen: Intracellular Genomic DNA Amplification for Downstream Applications in Sequencing and Sorting

**DOI:** 10.1101/2025.03.03.641299

**Authors:** P. Crossland, K. Schmidlin, I. Valli-Doherty, L. Brettner, K. Geiler-Samerotte

**Author notes:** co-first authors (P. Crossland, K. Schmidlin). co-senior authors (L. Brettner, K. Geiler-Samerotte).

## Abstract

Here, we introduce intracellular genomic amplification (INgen), a method that harnesses the cell membrane as a natural reaction chamber to amplify DNA within fixed, permeabilized cells. INgen employs a strand-displacing, isothermal polymerase together with biotinylated primers to achieve robust intracellular DNA amplification and efficient recovery of the amplified material for sequencing. This approach overcomes a long-standing barrier that has prevented important advances in single-cell and rare-cell genomics: the inability to amplify and recover sequenceable DNA from fixed cells. By pushing past this barrier, INgen provides a critical and previously inaccessible step toward scalable single-cell DNA sequencing without the need to isolate each cell into a separate reaction vessel. Using INgen, we demonstrate targeted and whole-genome amplification across diverse organisms, including *Saccharomyces cerevisiae*, *Bacillus subtilis*, and *Escherichia coli*. Together, these capabilities position INgen as a foundational advance that paves the way for the next generation of single cell sequencing methods including high-throughput single-cell sequencing without physical isolation, contamination-resistant amplification within intact cells, and rare-cell enrichment prior to genomic analysis.

## Introduction

Single-cell DNA sequencing has the potential to transform our ability to study genetic heterogeneity in mixed microbial communities (e.g., microbiomes) and in any population where cells are genetically diverse (e.g., tumors, drug-resistant infections, laboratory experiments, etc.). However, a major bottleneck for many single-cell ‘omics technologies is the dependence on physical cell isolation, which limits scalability and throughput. As a result, single-cell genomics studies are often limited to just hundreds (Gasch et al. 2017; Schmidlin et al. 2024; Venkataram et al. 2016) or even dozens of cells (Behringer et al. 2024; Zion et al. 2024). Additionally, isolation techniques often demand time-intensive labor, specialized expertise, and expensive instrumentation, further limiting their accessibility. Ironically, each cell’s membrane already naturally compartmentalizes its genetic material, providing a built-in reaction chamber. Yet, sequencing reactions are still carried out in separate vessels, a redundant and inefficient step given that the cell itself is perfectly isolated. This inherent feature suggests that using the cell as its own reaction chamber could significantly streamline single-cell genomic workflows.

Using the cell as its own reaction container has been extensively harnessed for single-cell transcriptomics (Brettner et al. 2024; Kuchina et al. 2021; Rosenberg et al. 2018; Cao et al. 2017; Martin et al. 2023; Blattman et al. 2020). However, intracellular DNA-based applications have not been widely implemented due to several technical challenges (Speel et al. 1999; Hully 1999), including the hurdle of amplifying DNA in formaldehyde-fixed cells in such a way that allows it to be sequenced. A robust method for intracellular DNA amplification that allows for downstream DNA sequencing thus stands to enable the optimization of higher-throughput single-cell DNA sequencing protocols.

A significant challenge in intracellular genomics arises from the need to fix cells with formaldehyde in order to turn them into sturdy reaction chambers. Fixing with formaldehyde stabilizes the cell and the genetic material inside, but the crosslinks that are generated are frequently cited as obstacles to efficient DNA amplification and can hinder sequencing (Kirsch et al. 2024; Clingenpeel et al. 2014; Pereira et al. 2022; O’Leary et al. 1994). While some studies suggest that formaldehyde can be reversed such that they do not severely inhibit amplification (Gavrilov and Razin 2009; Bagasra 2007), concerns about formaldehyde have driven efforts to perform FISH and FACS in live cells instead (Pereira et al. 2022). While short, engineered DNA sequences (15 bp) (Payne et al. 2021) and RNA transcripts have been successfully amplified in fixed cells and subsequently sequenced (Brettner et al. 2024; Rosenberg et al. 2018; Kuchina et al. 2021; Salatino et al. 2024), we found no reports of sequencing genomic DNA that was amplified inside of formaldehyde-fixed cells. Instead, recent intracellular methods randomly insert transposons that drive transcription of DNA to RNA, cleverly circumventing the challenge of intracellularly amplifying and sequencing DNA (Vitak et al. 2017; Yin et al. 2019).

To address this challenges, we introduce intracellular genomics (INgen), a novel method that enables intracellular DNA amplification and subsequent DNA sequencing. By overcoming long-standing barriers, this advance has the potential to unlock important downstream applications such as higher throughput single-cell DNA sequencing without the need for cell isolation. In this study, we report the methodological details of INgen, including the key innovations that explain why it works. We then demonstrate successful sequencing of target genes up to 3kb and whole genomes in diverse organisms, including *S. cerevisiae, B. subtilis,* and *E. coli*. We also report control experiments confirming that the sequenced DNA originates from intracellular amplification, rather than leaked or native genomic DNA. We show that up to 14% of the yeast genome can be amplified within a single cell. Finally, we demonstrate successful fluorescence-activated cell sorting and sequencing of cells containing intracellularly amplified DNA.

In sum, INgen is a robust and versatile method for intracellular DNA amplification in diverse cell types, paving the way for a wide array of applications, including high-throughput single-cell sequencing, genotype-based cell sorting, and rare cell genomics. By breaking this critical barrier of amplifying and sequencing DNA from fixed cells, INgen stands to enable advances in single-cell ‘omics with broad potential applications in microbial ecology, cancer genomics, and evolutionary biology.

## Results

### A flexible method for intracellular DNA Amplification (INgen)

In this manuscript, we address the challenge of amplifying DNA within cells and sequencing this DNA. One challenge associated with intracellular DNA amplification described in the literature is that the temperature cycling required for polymerase chain reaction can weaken cell membranes and cause DNA leakage (Speel et al. 1999; Hully 1999). Another challenge described in the literature is that the formaldehyde cross-links that turn fixed cells into sturdy reaction chambers come with a negative side effect: they block enzymes such as DNA polymerase from accessing DNA (Kirsch et al. 2024; Clingenpeel et al. 2014; Pereira et al. 2022; O’Leary et al. 1994). Our use of Phi29 polymerase, a commonly used enzyme for amplifying DNA, though not typically employed intracellularly (Lasken 2007; Nelson 2014), contributes to our success in overcoming these challenges. Previous work suggests that phi29 polymerase can overcome formaldehyde crosslinks through its robust strand-displacement activity (Lee et al. 2006; Aviel-Ronen et al. 2006; Tang et al. 2020). Furthermore, Phi29 is an isothermal polymerase, meaning it does not require the thermal cycling that has been associated with membrane disruption, increased risk of extracellular amplification, and false signals (Speel et al. 1999). Instead, our protocol includes only a short, controlled pre-incubation at 72°C, prior to polymerase addition, to facilitate primer annealing.

Another way the INgen protocol addresses the challenge of recovering DNA from formaldehyde-fixed cells is through biotinylated primers. These enable the selective capture of intracellularly amplified DNA on magnetic streptavidin beads following cell lysis. Streptavidin-bound DNA can be washed multiple times, allowing us to remove harsh reagents used during the lysis of fixed cells. This purification step helps ensure that subsequent amplification reactions, such as those used for preparing DNA for sequencing, can proceed efficiently.

The INgen workflow consists of four key steps: (1) fixation and permeabilization of cells to stabilize the cellular structure and allow reagent access (**Fig 1a–c**), (2) intracellular DNA amplification using an isothermal polymerase (**Fig 1d**) and biotinylated primers, (3) cell washing, cell lysis, and isolation of newly synthesized DNA via streptavidin-coated beads (**Fig 1e–f**), and (4) downstream sequencing of amplified genomic material. We demonstrate that intracellular amplification and subsequent sequencing of the amplified products is not only feasible but also effective across multiple contexts, underscoring the utility of this method for downstream single-cell sequencing applications.

**Figure 1.**
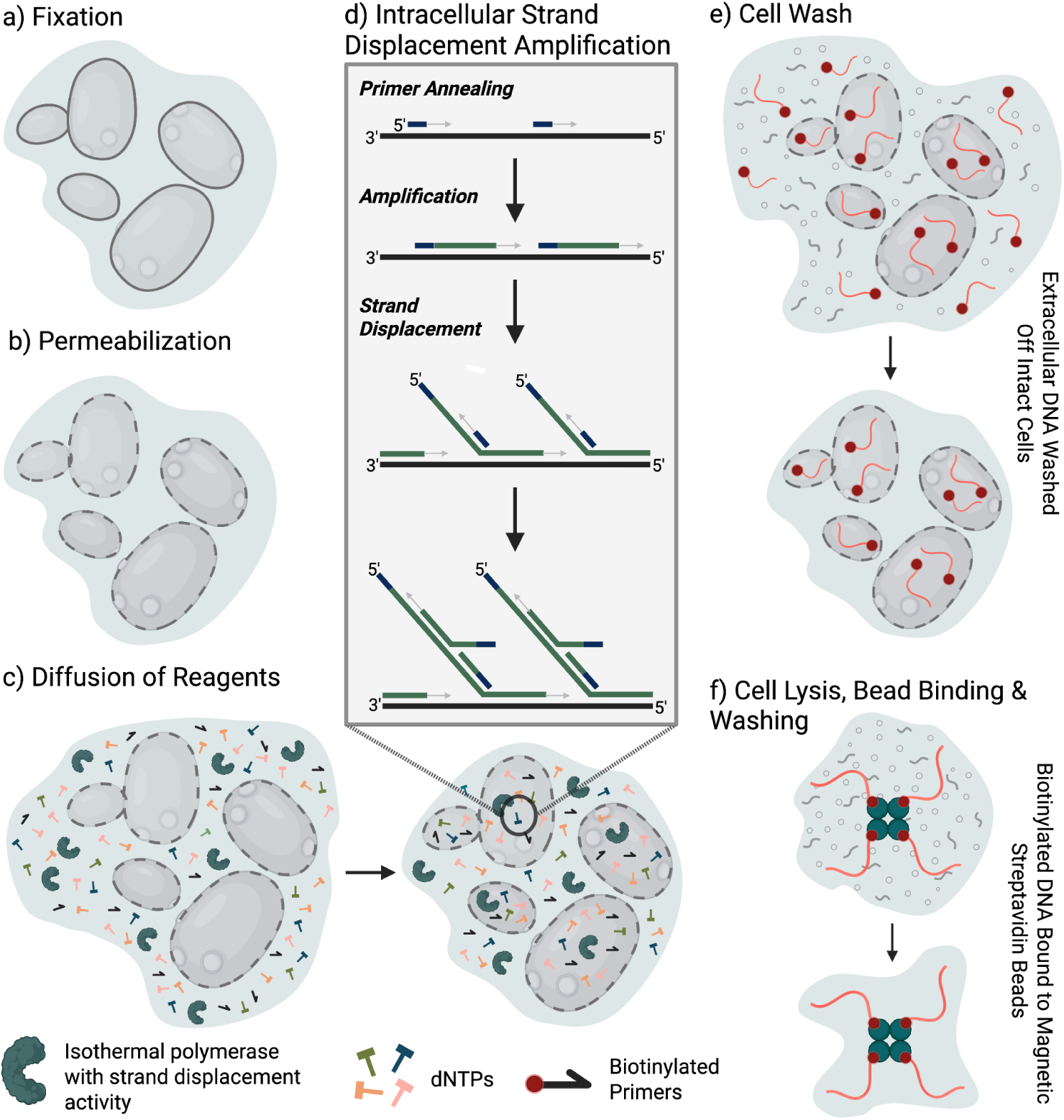
Schematic of the intracellular DNA amplification (INgen) workflow. This robust and flexible protocol can be altered at each step based on the cell type being amplified. **a)** Fixation stabilizes the cell, which needs to remain intact during the entire amplification reaction. It varies by organism: 4% formaldehyde for yeast and 70% ethanol for bacteria. **b)** Permeabilization allows reagents to diffuse inside cells. Yeast are treated with zymolyase and triton-X, while bacterial membranes become permeable during ethanol fixation. **c)** Reaction reagents, including phi29, dNTPs, and primers, are diffused into the cell by the crowding agent sorbitol. See methods for the detailed master mix recipe. **d)** Intracellular strand displacement amplification (ISDA) proceeds isothermally via primer annealing, polymerase extension, and strand displacement, generating concatemeric DNA products within each intact cell (Spits et al. 2006). **e)** Cell washing removes extracellular DNA and unincorporated reagents while maintaining the integrity of amplified cells. **f)** Cell lysis releases amplified DNA, which contains incorporated biotin. The biotinylated DNA is bound to magnetic streptavidin-coated beads, enabling subsequent purification and downstream processing such as sequencing library preparation.

To assess the applicability of our method across taxa, we tested intracellular amplification in the eukaryotic model organism *Saccharomyces cerevisiae* (yeast) (**Fig. 2 & 4**) and two prokaryotic species, *Bacillus subtilis* (a gram-positive bacterium) (**Fig. S6**) and *Escherichia coli* (a gram-negative bacterium) (**Fig. 2 & 3**). Using species-specific fixation and permeabilization protocols, we successfully amplified genomic material in all organisms. For yeast, fixation with formaldehyde followed by treatment with zymolyase and Triton-X enabled permeabilization, while ethanol fixation alone was sufficient for *E. coli* and *B. subtilis*, and did not require additional permeabilization steps. These results demonstrate that our platform is robust to the distinct cell wall and membrane compositions of Gram-negative and Gram-positive bacteria, as well as fungi, making it broadly applicable across phylogenetically diverse organisms.

**Figure 2.**
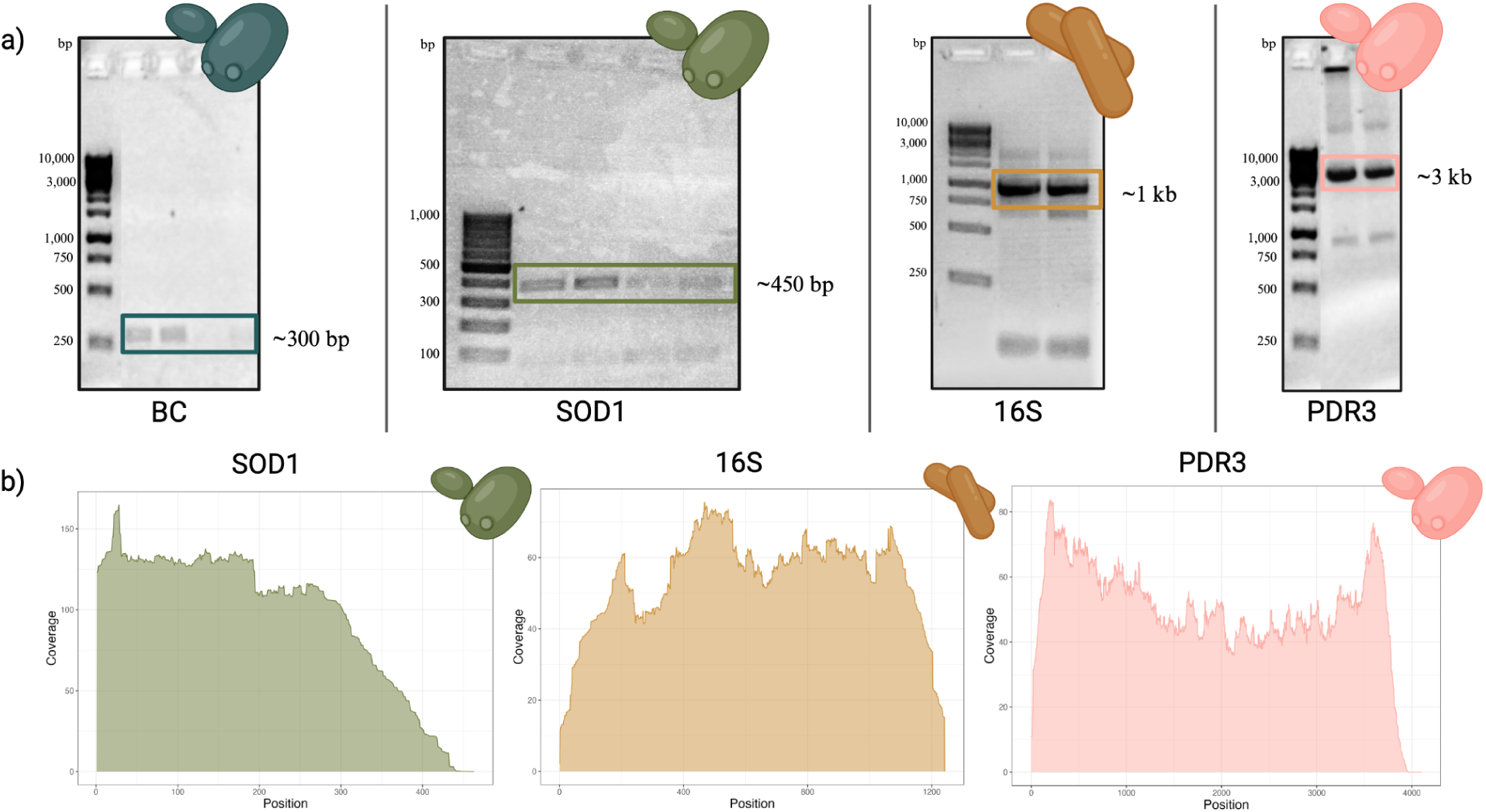
Targeted amplification and sequencing of specific genomic regions. **a)** Gel electrophoresis of amplified DNA fragments targeting different genomic loci. BC: barcode region (∼300 bp, teal), *SOD1* (∼450 bp, green), 16S rRNA (∼1 kb, orange), and *PDR3* (∼3 kb, pink). Bands corresponding to the expected fragment sizes are highlighted with colored boxes. **b)** Sequencing coverage plots for *SOD1* (green), 16S rRNA (orange), and *PDR3* (pink) show the read depth across each amplified region. Coverage varies across each fragment, but most bases are covered over 15X. Because amplification is carried out using phi29 polymerase with gene-specific primers in a strand-displacing, isothermal reaction, amplification proceeds unidirectionally from the primer rather than symmetrically from both ends. This results in non-uniform coverage skewed in the direction of polymerase extension. Additional variation may arise from differences in primer annealing efficiency or DNA accessibility within fixed cells.

**Figure 3.**
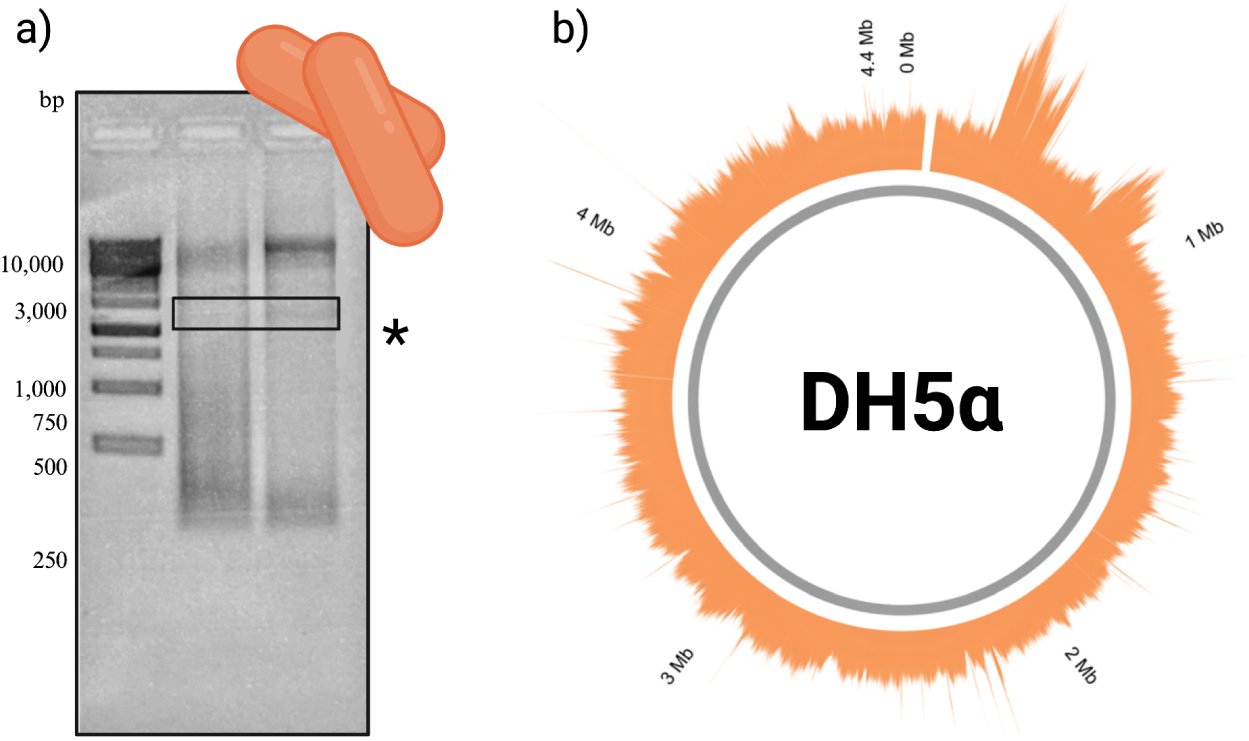
Whole-genome amplification and sequencing of *E. coli* DH5α. a) Agarose gel electrophoresis of amplified *E. coli* DNA showing a high-molecular-weight smear indicative of successful whole-genome amplification. The boxed region highlights a representative fragment size within the amplification range. The asterisk (*) marks an additional band corresponding to the expected size of a 16S rRNA amplicon, as 16S primers were included alongside random hexamers in the same reaction, demonstrating that multiple primer types can be used concurrently. b) Sequencing coverage across the *E. coli* DH5α genome. Read depth is shown in orange, coverage exceeds 30x across 99.4% the 4.6-Mb circular chromosome.

**Figure 4.**
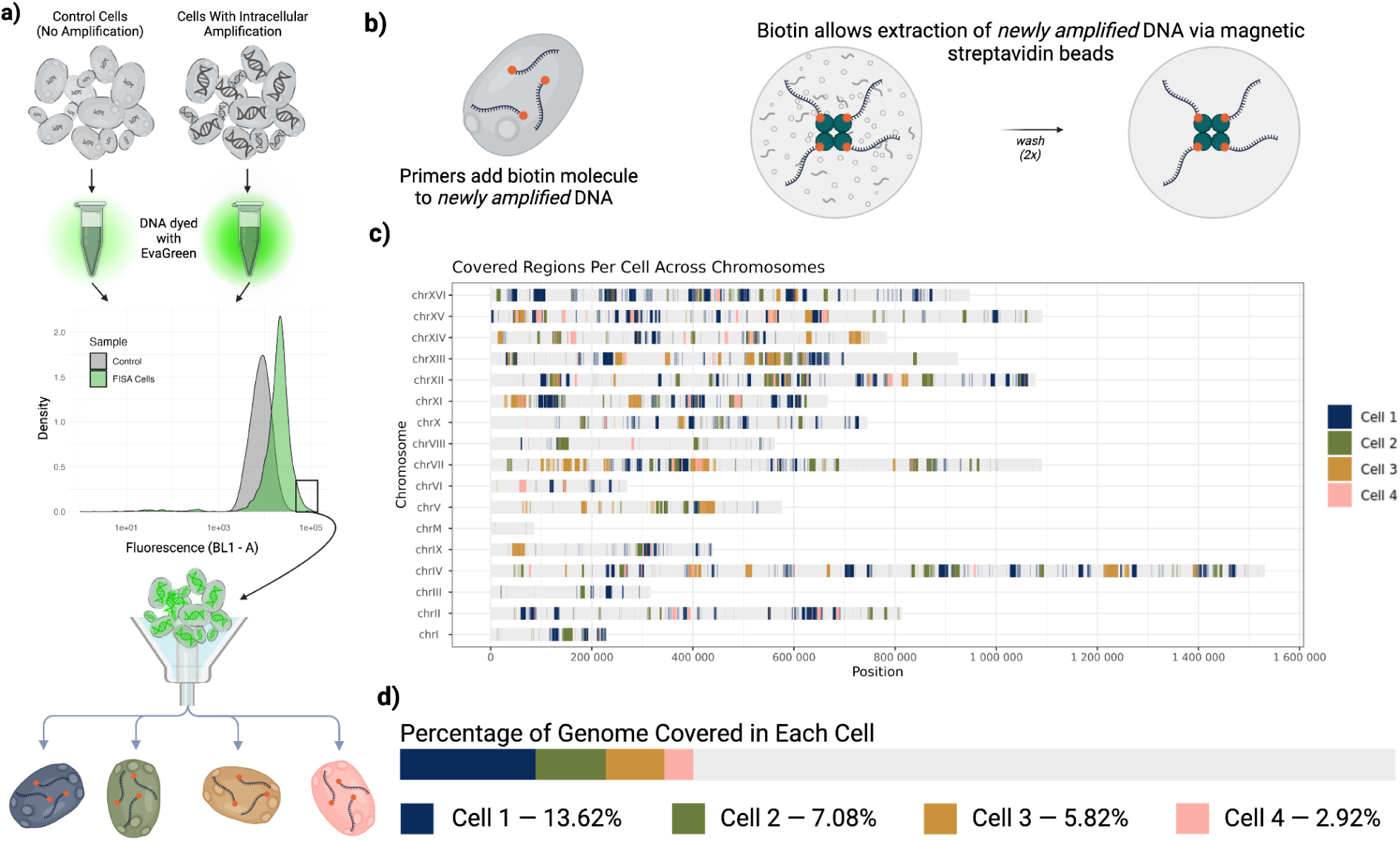
Intracellular Genome Amplification and Sequencing Coverage Analysis. (a) Schematic representation of intracellular genome amplification. Control cells (left) do not undergo amplification, while treated cells (right) contain newly synthesized DNA. DNA was stained with EvaGreen, and fluorescence intensity was measured by flow cytometry, revealing increased fluorescence in cells that underwent intracellular amplification compared to controls. Amplified cells are then sorted based on high fluorescent signals for sequencing. (b) Biotinylated primers enable the selective extraction of newly amplified DNA using magnetic streptavidin beads, followed by multiple washes to purify the sample. (c) Genome coverage analysis of four individual cells mapped to *S. cerevisiae* chromosomes. Each colored segment represents a covered region for a specific cell. (d) Summary of genome coverage for each sequenced cell. The percentage of the genome covered is highest in Cell 1 (13.62%) and lowest in Cell 4 (2.92%).

### Intracellular amplification of targeted regions and recovery of DNA for sequencing

To evaluate the ability of INgen to selectively amplify specific genomic regions, we designed biotinylated primers targeting genes of varying lengths, including three genes from yeast (SOD1, 450bp; PDR3, 3,000bp; and an engineered barcode BC, 300bp) and one gene from bacteria (16S rRNA, 1,000bp). For each reaction, we performed the INgen protocol on roughly 1 million cells of the target cell type following the method described above (**Fig. 1**). Biotinylated primers were annealed to DNA during a 5-minute incubation at 72 °C, performed after all reaction components, except Phi29 polymerase, had been diffused into the cells. Following this step, Phi29 polymerase was added, and cells were incubated for 16 hours at 30 °C. After incubation, cells were lysed, and the newly synthesized, biotin-labeled DNA was isolated using streptavidin-coated beads. Once bound and washed, a secondary ‘off-the-beads’ amplification reaction using targeted primers was performed to generate sufficient DNA for library preparation, which prepares DNA for next-generation sequencing.

Gel electrophoresis analysis confirmed successful targeted amplification off-the-beads, with distinct bands of the expected sizes observed for all tested loci (**Fig. 2A**). While we didn’t attempt to amplify any regions above 3kb, the highly processive nature of phi29 (Blanco et al. 1989) and our own whole genome data (**Figure S3, S4, S5)** suggests that longer targets could also be amplified. Following size confirmation on an agarose gel, DNA fragments were prepared for sequencing using the Illumina DNA Flex kit. Targeted genomic regions were successfully sequenced to a minimum depth of 15X (**Fig. 2B**). The ability to target multiple genes across multiple species highlights the flexibility and broad applicability of this method.

To confirm that DNA amplification is happening intracellularly, rather than extracellularly on leaked DNA, we performed a control experiment where we spiked varying amounts of bacterial DNA into a reaction vessel containing yeast cells (**Fig. S1A**). We performed the INgen protocol on these mixtures using 16S primers targeting the bacterial DNA. After the 16-hour amplification step, we removed the supernatant and used the same 16S primers to reamplify 16S bacterial DNA, obtaining clear bands on an agarose gel (**Fig. S1A;** pre-wash). However, after washing the cells twice, PCR reactions using subsequent supernatants did not produce robust bands (**Fig. S1A;** wash 2). Given that we spiked in more DNA than we would ever expect to be present extracellularly, and still could not achieve robust amplification after wash steps, we conclude that the amplicons produced during a typical INgen reaction (**Fig. 2**) are not predominantly created extracellularly by priming and elongating DNA fragments that leaked out of cells.

To confirm that the amplicons in **Figure 2** do not reflect amplification of native genomic DNA post-cell lysis, we performed a second control experiment (**Fig. S1B**). Native genomic DNA should be washed away post-cell lysis, leaving only streptavidin-bound products of intracellular amplification. To test this, we spiked large amounts of non-bead-bound bacterial 16S DNA into reactions post-cell-lysis. Amplification reactions performed using the supernatant from subsequent wash steps failed to produce robust 16S products on an agarose gel after just two washes (**Fig. S1B**). We also performed a control experiment to confirm that non-bead-bound DNA cannot associate with streptavidin beads (**Fig. S2**). These control experiments support our conclusion that amplification is specific to intracellularly amplified, biotinylated DNA and not native genomic DNA that fails to be washed away. (Also see **Fig. 4**, where we confirm intracellular amplification by showing cells that underwent INgen have a stronger fluorescent signal after DNA stain).

While intracellular DNA amplification has been attempted before, prior methods produced limited or unusable material that could not be sequenced (Speel et al. 1999; Pereira et al. 2022). INgen overcomes this barrier by enabling robust intracellular amplification and recovery of sequenceable DNA across diverse cell types and genomic targets. Multiple controls confirm that amplification occurs within cells rather than from leaked or native DNA. By making it possible to sequence DNA amplified inside fixed cells, INgen has the potential to transform what was previously a technical dead end into a versatile platform for single-cell genomics.

### Whole genome amplification with random hexamers

Beyond targeted genes, we next asked whether INgen could be extended to amplify and sequence entire genomes. To test this, we performed INgen using biotinylated random hexamers in ∼1 million *E. coli* cells. Following intracellular amplification, we isolated the newly synthesized, biotin-labeled DNA, re-amplified it off the beads using random hexamers, and sequenced the resulting DNA pool.

Gel electrophoresis of the off-the-beads PCR product revealed smears spanning a broad range of fragment sizes, consistent with successful whole-genome amplification. Fragment sizes ranged from 200 bp to 50 kb (**Fig. 3A**). TapeStation analysis further confirmed that intracellular amplification generated long DNA fragments, with an average size of ∼20kB, and some spanning up to several hundred kilobases (**Fig. S3, S4 & S5**), consistent with the expected performance of phi29 polymerase. These results suggest that formaldehyde crosslinking does not significantly inhibit this polymerase’s activity in our reaction conditions.

The collective reads from many INgen-amplified cells reconstructed the entire *E. coli* genome (**Fig. 3B**). From a single intracellular PCR reaction containing approximately one million cells, 99.4% of the *E. coli* genome was covered at a depth greater than 30× (**Fig. 3B**). These results demonstrate that INgen supports robust amplification across diverse genomic loci within fixed cells, expanding its utility beyond targeted assays and setting the stage for comprehensive, isolation-free, single-cell genome sequencing, assuming each cell’s DNA could be barcoded in a similar way as is done for RNA (Brettner et al. 2024; Kuchina et al. 2021; Rosenberg et al. 2018; Vitak et al. 2017).

The INgen method could also be deployed to address a persistent challenge in traditional isolation-based single-cell whole-genome sequencing: contamination. Because isolation-based scDNAseq experiments begin with extremely limited DNA (that from a single isolated cell), even trace amounts of exogenous DNA can be amplified along with the genome of interest.

Random hexamers exacerbate this problem since they bind indiscriminately to any DNA, including contaminants introduced from reagents, tubes, or gloves (Nawy 2014; Lusk 2014; Lux et al. 2015). INgen mitigates this issue by using the cell itself as a sealed reaction vessel, thereby physically restricting amplification to the DNA already present within that cell. In addition, the use of biotinylated primers enables selective recovery of intracellularly amplified DNA, while wash steps remove non-biotinylated or extracellularly biotinylated DNA (**Fig. S1 & S2**). Together, these features could minimize contamination and improve the accuracy and reliability of single-cell genomic sequencing.

### Successful sorting and sequencing of individual cells with intracellularly amplified DNA

To further validate that DNA amplification is occurring intracellularly, we designed control experiments in yeast cells where we omitted either polymerase or primers from our intracellular reactions. We then stained control and test yeast cells with EvaGreen, a fluorescent dye that binds to all double-stranded DNA, and compared fluorescence distributions. Indeed, test populations exhibited significantly greater average fluorescence compared to controls because the test reactions possessed all necessary reagents to intracellularly amplify DNA (**Fig. 4A**). This provides further evidence that amplification is occurring intracellularly.

We next used flow cytometry to sort and select test cells exhibiting brighter fluorescence than measured in any control cell (**Fig 4A**). The fluorescent cells that met our criteria were sorted into individual vessels so that we could confirm they each contain intracellularly amplified DNA that is viable for sequencing. Our primary objective was to determine if the brightest cells indeed contained sequenceable, biotinylated DNA. Our second objective was to learn what fraction of the genome was amplified within the brightest cells.

We processed 4 single cells. Each cell was lysed using Proteinase K, and the newly synthesized, biotinylated DNA was captured with streptavidin-coated beads and washed several times to remove native genomic DNA (**Fig. 4B**). For all four cells, the bead-bound DNA was then re-amplified using random hexamers, which generated enough starting material (>5 ng/uL) to construct sequencing libraries. This suggests all four cells contained biotinylated DNA amplicons.

Next, we asked how much of the genome we had amplified in any one cell. Reported single-cell whole-genome recovery yields are as high as ∼80–90 % in mammalian systems (Kojima et al. 2021; Leung et al. 2016). However, in challenging contexts, such as microbial or fixed cells, recovery rates can be an order of magnitude lower (de Bourcy et al. 2014; Zong et al. 2012). In such systems, researchers often reconstruct complete genomes computationally by clustering partial genome fragments across multiple cells or bins (López-Escardó et al. 2017; Andresen et al. 2021). Indeed, recent advances in computational methods can group cells from the same species and accurately rebuild full genomes even from ultra-low-coverage single-cell data, sometimes as little as 0.03× per cell (Rozhoňová et al. 2022; Khan and Mallory 2023; Menon 2019). Similar strategies have also been used to resolve clonal architectures in tumors and to reconstruct protist genomes from sparse or metagenomic data (Dia and Cheeseman 2021; Wideman et al. 2020; Sieracki et al. 2019).

In terms of assembling genomes by clustering partial reads from many cells, a key strength of INgen is that intracellular amplification can be performed in parallel across thousands of pooled cells. Although INgen does not yet incorporate barcoding or cell-specific identifiers, in principle, INgen could be paired with combinatorial indexing approaches that have already transformed single-cell RNA sequencing (Brettner et al. 2024; Kuchina et al. 2021; Rosenberg et al. 2018; Vitak et al. 2017). Such coupling could enable genome recovery by clustering partial, cell-labeled genomes, potentially enhanced by primers that target anchor regions such as 16S rRNA genes. Even a few percent of a single cell’s genome may therefore be sufficient for this kind of analysis, as demonstrated in low-coverage single-cell RNA-seq studies where sparse transcriptome coverage still resolves cell-type relationships (Brettner et al. 2024; Kuchina et al. 2021; Pollen et al. 2014).

At this stage, our goal for INgen is not to optimize genome recovery, but simply to benchmark performance. Because DNA recovery from formaldehyde-fixed cells has historically been so difficult, our first aim was to test whether we could obtain enough amplified DNA to make any downstream analyses conceivable. Given that we selected from the cells with the strongest fluorescent signal, this experiment will not report the average amount of intracellular amplification per cell, but the upper limit, given the extent of optimization thus far. Our sequencing data revealed that between 3 and 13% of the yeast genome was recovered from each of the four cells we processed. There was minimal overlap between the genomic regions recovered from different cells (**Fig. 4C**), such that combining reads from all four cells resulted in the recovery of 29.44% of the yeast genome. This high level of per-cell coverage demonstrates that INgen is capable of generating sufficient information for accurate genome reconstruction via cell clustering and thus provides a path toward high-throughput single-cell DNA sequencing.

In sum, INgen establishes a versatile foundation for next-generation single-cell genomics. By enabling sequencing of DNA from formaldehyde-fixed cells after FACS, it could transform rare-cell genomics, allowing researchers to sort desired genotypes based on fluorescence and then sequence the enriched genotypes within the sorted pool. INgen can also complement existing isolation-based scDNAseq approaches by pre-amplifying intracellular DNA to boost signal and reduce contamination. Most excitingly, INgen opens the door to high-throughput single-cell DNA sequencing without physical isolation, since combinatorial barcoding strategies already proven for RNA can now, in principle, be applied to DNA. By overcoming the key barrier to amplifying and sequencing DNA inside fixed cells, INgen sets the stage for scalable, pooled single-cell genomics across diverse biological systems.

## Discussion

INgen breaks a decades-old barrier: it enables robust amplification and recovery of DNA from within fixed cells. This is a prerequisite step that has long prevented scalable single-cell DNA sequencing. INgen overcomes major challenges that previously made intracellular DNA amplification and sequencing impossible. Using phi29 polymerase, a highly processive, strand-displacing enzyme, we achieved strong intracellular amplification under isothermal conditions, avoiding the high-temperature cycles that damage membranes. Incorporating biotinylated primers allowed selective capture of newly synthesized DNA after lysis, enabling downstream purification and sequencing.

Across all tested conditions, different fixation and permeabilization methods, species (yeast, *E. coli*, and *Bacillus subtilis*), and amplification targets, INgen performed robustly. We validated targeted amplification and sequencing of specific genomic loci and demonstrated successful whole-genome amplification in bacteria, recovering over 99% of the *E. coli* genome from a pooled intracellular reaction. We further showed that individual yeast cells can yield up to 13% genome recovery, a level sufficient to enable downstream clustering or reconstruction when applied across many cells. By demonstrating effective intracellular DNA amplification in both prokaryotes and eukaryotes, INgen establishes a generalizable foundation for scalable single-cell DNA analysis.

The ability to amplify DNA inside fixed cells without compromising membranes or sequenceability transforms what can now be attempted in single-cell genomics. INgen’s design, amplifying DNA intracellularly, labeling it via biotinylation, and then purifying it for sequencing, provides a modular framework for next-generation developments, including combinatorial barcoding, genotype-based cell sorting, and high-throughput analysis of complex populations such as microbiomes or tumors.

### Comparison with Existing Amplification and Sequencing Techniques

INgen is more versatile than other intracellular amplification methods, such as in situ PCR (IS-PCR), because INgen allows sequencing of the amplified material. IS-PCR instead has traditionally been used to localize cells containing particular genetic elements within tissues or tumors (Hully 1999; Murray 1993; Bagasra 2007; Janiszewska et al. 2015) by enhancing the signal for probe hybridization (Muro-Cacho 1997; Speel et al. 1999; Hully 1999; Nuovo 1997). Specific applications of IS-PCR have included detecting HIV (Bagasra et al. 1992; Gibellini et al. 1998) and malaria in human cells (Hashimoto et al. 2018), analyzing oncogenes in tumors (Janiszewska et al. 2015; Salatino et al. 2024), and detecting pathogens in environmental samples (Tani et al. 1998). However, the widespread adoption of IS-PCR has been stalled by technical issues, including compromised cell membranes, extracellular amplification, non-uniform amplification, and false signals (Speel et al. 1999; Yin et al. 2019; O’Leary et al. 1996). Further, IS-PCR is usually performed on paraffin-embedded samples, which restricts downstream applications such as sorting labeled cells for ‘omics analyses of genotypes of interest (Hashimoto et al. 2018; Janiszewska et al. 2015; Bagasra 2007). INGen overcomes many of the limitations of IS-PCR because it works with cells in solution, amplifies DNA at low temperatures that do not result in compromised membranes, and biotinylates the amplicons which eliminates false signals from extracellularly amplified DNA.

Compared to conventional multiple-displacement amplification (MDA) methods, INgen also poses some advantages. MDA requires live-cell isolation and lysis, which limits scalability and complicates work with fragile or fixed samples (Spits et al. 2006). INgen overcomes this limitation by performing amplification prior to lysis, allowing DNA to be recovered even from preserved or formaldehyde-fixed cells. This feature expands access to clinical, environmental, and archival samples that cannot easily be processed by existing single-cell techniques (Huang et al. 2015; Shi et al. 2004). However, INgen may be sensitive to other limitations related to phi29 sequencing, including a bias towards amplifying plasmids.

Finally, while single-cell RNA-seq revolutionized transcriptomic analysis, it cannot directly capture genomic variation (Brettner et al. 2024; Kuchina et al. 2021; Rosenberg et al. 2018). INgen complements such methods by enabling intracellular amplification of genomic DNA, paving the way for the detection of non-coding mutations, structural variants, and other lineage-defining changes that are invisible to RNA-based assays. Although INgen currently focuses on intracellular DNA amplification and recovery rather than full barcoding or multiplexed sequencing, breaking this technical barrier establishes the essential groundwork for those next steps.

In sum, INgen transforms what was once a dead end, the inability to amplify and recover DNA from fixed cells, into a versatile platform for next-generation single-cell genomics. Continued optimization, such as incorporating newer variants of phi29 polymerase, could further enhance its performance. By overcoming a fundamental barrier, INgen opens new possibilities for scalable, pooled, and barcoded single-cell DNA sequencing across a broad range of biological systems.

## Methods

### Fixation and Permeabilization

Fixation and permeabilization are crucial steps in our method, as they maintain cellular integrity while enabling reagent diffusion for intracellular genome amplification. This balance is necessary to preserve cell structures while ensuring effective reagent penetration. To achieve this, we evaluated multiple fixation strategies to identify optimal conditions.

Formaldehyde-based fixation provided a robust approach by crosslinking proteins and stabilizing cellular architecture while preserving nucleic acids. We tested protocols ranging from short incubations at room temperature (e.g., 10 minutes, as described in Payne et al., 2021) to extended fixation at 4°C, a commonly used method for yeast preservation (cite). Both protocols maintained cell morphology and supported efficient downstream amplification.

Ethanol fixation/permeabilization offered an alternative strategy. Ethanol rapidly precipitates proteins and disrupts membranes, enhancing permeability (cite). While harsher than formaldehyde, this method effectively permeabilized bacterial cells such as E. coli DH5α. Its simplicity and rapid processing time (under 15 minutes) made it an attractive option.

Interestingly, the fixation method had minimal impact on genomic amplification efficiency or fluorescent labeling, underscoring the robustness of our approach. However, differences in cell durability and fluorescence intensity were observed, suggesting that formaldehyde fixation may better preserve intracellular targets for flow cytometry.

Following fixation, permeabilization was required to facilitate reagent entry. We tested enzymatic treatments, such as zymolyase digestion for yeast cell walls, and detergent-based methods using Tween-20 and Triton X-100. Zymolyase treatment efficiently converted yeast cells into spheroplasts while maintaining structural integrity. Detergent-based approaches provided a rapid, scalable alternative applicable to both yeast and bacterial cells.

These optimizations of fixation and permeabilization enabled intracellular amplification across diverse cell types. Our approach refines existing protocols to meet the specific demands of fluorescently labeled intracellular genome amplification.

### Intracellular Amplification

The central innovation of our method is the ability to amplify genomic DNA directly within fixed and permeabilized cells. By performing amplification intracellularly, we mitigate biases inherent in traditional single-cell sequencing workflows, such as uneven amplification or the loss of fragile genomic regions during extraction. Instead, our method leverages the cellular environment itself as a natural reaction vessel, enabling robust, contained, and targeted DNA synthesis.

The selection of an appropriate DNA polymerase was critical to the success of this approach. We employed phi29 DNA polymerase, a highly processive, strand-displacing enzyme optimized for isothermal amplification and capable of functioning despite the presence of residual fixative or membrane components. This enzyme supported consistent amplification across a variety of cell types, including yeast and bacteria.

The heart of our method is the controlled intracellular reaction setup. Fixed and permeabilized cells are first incubated with a reaction mixture containing 1X phi29 buffer (New England Biosystems), 0.2 mg/mL BSA, 250 µM dNTPs, 1.15 M sorbitol, and 2 µM biotinylated primers (either gene-specific or random hexamers). Approximately 5 µL of a 1 × 10⁶ cells/mL suspension (∼5 × 10³ cells total) is added to 45 µL of reaction mixture to yield a final volume of 50 µL. The mixture is pre-incubated at 72 °C for 5 minutes to facilitate primer annealing while preserving cell structure. Phi29 polymerase (New England Biosystems) is then added (final concentration: 100 U per 50 µL reaction), and the complete reaction is incubated at 30 °C for 16 hours under isothermal conditions.

By avoiding traditional thermal cycling, INgen reduces membrane disruption, minimizes nonspecific extracellular amplification, and decreases the risk of amplification artifacts or false-positive signals. Amplification products remain confined within the cell and covalently bound to biotinylated primers, enabling their selective purification using streptavidin-coated magnetic beads. This approach preserves genomic integrity, supports uniform genome-wide amplification, and improves the detection of low-abundance genetic features such as rare mutations or transposable elements.

### Identifying Highly Amplified Cells: EvaGreen Staining for Genome Quantification and Sorting

To quantify intracellular genome amplification, cells were stained with EvaGreen, a DNA-binding dye that fluoresces upon binding to double-stranded DNA. Cells were incubated with EvaGreen for 30 minutes to ensure sufficient dye uptake while minimizing background fluorescence. Following incubation, excess dye was removed by centrifugation, and the cells were washed to prevent non-specific staining.

Fluorescence intensity was measured using an Attune flow cytometer, allowing for precise quantification of amplified DNA within individual cells. Cells exhibiting high fluorescence—indicative of robust genome amplification—were identified and selected for sorting. Using fluorescence-activated cell sorting (FACS), highly amplified cells were isolated and deposited into individual wells of a 96-well plate for downstream processing. This approach enabled the targeted recovery of cells with successful intracellular amplification, improving the efficiency and accuracy of genomic analysis.

### DNA Recovery and Sequencing Preparation: Capturing and Amplifying Biotinylated Genomes

Following intracellular amplification, cells were lysed using proteinase K to release genomic material. To inactivate the proteinase K, 5 μL of 100 μM PMSF (resuspended in isopropanol) was added to each tube, followed by a 10-minute incubation at room temperature. No high temperature step was included after lysis. Genomic DNA within fixed cells remains largely insoluble under our lysis conditions (non-ionic detergent, Proteinase K), and is not expected to be released into the supernatant unless the cells are physically or enzymatically disrupted.

For selective capture of intracellularly amplified, biotinylated DNA, 100 μL of resuspended streptavidin-coated C1 magnetic beads were added to each sample and incubated at room temperature for 60 minutes with agitation. The samples were then placed on a magnetic rack, and the supernatant was removed. Beads were resuspended in 250 μL of 1X wash solution, agitated for 5 minutes, and subjected to two consecutive wash steps. After the final wash, beads were resuspended in 250 μL of 10 mM Tris-HCl (pH 8.0) with 0.1% Tween-20. At this stage, beads could either be rinsed with molecular-grade water and processed immediately or stored overnight at 4°C in a Tris-Tween buffer.

Amplification off-the-beads was performed by adding bead bound DNA to a 50 µl reaction mix which contained 1X phi29 buffer, 0.2 mg/mL BSA, 250 µM dNTPs, 1.15 M sorbitol, and 2 µM random hexamers. Libraries were constructed using the Genomic DNA by ligation kit (SQK-LSK109) following manufacturer’s instructions. Libraries were quantified using Qubit and sequencing was performed either on MinION using a FLO-MIN106D flow cell, or was prepared for Illumina sequencing, using the Illumina DNA Prep library preparation kit.

### Example Fixation and Permeabilization Permutations

#### a) Overnight Formaldehyde Fixation with Zymolyase and Tween-20 Permeabilization

At the time of sampling, 3 mL of yeast cultures, grown to saturation in YPD medium, were immediately spun down in a room-temperature centrifuge at 5000 g for 3 minutes. The cell pellet was fixed with 4% formaldehyde and incubated overnight at +4°C on a shaker. Fixed cells were centrifuged at 5000 g for 3 minutes, the supernatant was discarded, and the pellet was resuspended in ice-cold Buffer B (1.2 M sorbitol, 0.1 M potassium phosphate dibasic, pH 7.5). Zymolyase (1 mg/mL) was added, and cells were incubated at 30°C for 15 minutes. Permeabilization was performed with 0.1% Tween-20 at room temperature for 10 minutes before proceeding to downstream steps.

#### b) Overnight Formaldehyde Fixation with Zymolyase and TritonX-100 Permeabilization

At the time of sampling, 3 mL of yeast cultures, grown to saturation in YPD medium, were immediately spun down in a room-temperature centrifuge at 5000 g for 3 minutes. The cell pellet was fixed with 4% formaldehyde and incubated overnight at +4°C on a shaker. Fixed cells were centrifuged at 5000 g for 3 minutes, the supernatant was discarded, and the pellet was resuspended in ice-cold Buffer B (1.2 M sorbitol, 0.1 M potassium phosphate dibasic, pH 7.5). Zymolyase (1 mg/mL) was added, and cells were incubated at 30°C for 15 minutes. Permeabilization was performed with 0.1% TritonX-100 at room temperature for 10 minutes before proceeding to downstream steps.

#### c) 10 Minute Formaldehyde Fixation with Zymolyase and TritonX-100 Permeabilization (“INgen fix”)

At the time of sampling, 3 mL of yeast cultures, grown to saturation in YPD medium, were immediately spun down in a room-temperature centrifuge at 5000 g for 3 minutes. The cell pellet was fixed with 4% formaldehyde for 10 minutes at room temperature (+20°C). Fixed cells were centrifuged at 5000 g for 3 minutes, the supernatant was discarded, and the pellet was resuspended in ice-cold Buffer B (1.2 M sorbitol, 0.1 M potassium phosphate dibasic, pH 7.5). Zymolyase (1 mg/mL) was added, and cells were incubated at 30°C for 15 minutes. Permeabilization was performed with 0.1% TritonX-100 at room temperature for 10 minutes before proceeding to downstream steps.

#### d) 30 Minute Formaldehyde Fixation with Zymolyase and Overnight Ethanol Permeabilization (“FISH fix”)

At the time of sampling, 3 mL of yeast cultures, grown to saturation in YPD medium, were immediately spun down in a room-temperature centrifuge at 5000 g for 3 minutes. The cell pellet was fixed with 4% formaldehyde for 30 minutes at room temperature (+20°C) and then transferred to +4°C for overnight fixation on a shaker. Fixed cells were centrifuged at 5000 g for 3 minutes, the supernatant was discarded, and the pellet was resuspended in ice-cold Buffer B (1.2 M sorbitol, 0.1 M potassium phosphate dibasic, pH 7.5). Zymolyase (1 mg/mL) was added, and cells were incubated at 30°C for 15 minutes. The permeabilization step used ethanol treatment overnight at room temperature before downstream steps.

#### e) Ethanol Fixation and Permeabilization

At the time of sampling, 3 mL of *Escherichia coli* cultures, grown to the desired density, were immediately spun down in a room-temperature centrifuge at 5000 g for 3 minutes. The cell pellet was resuspended in 70% ethanol and incubated at room temperature (+20°C) for 10 minutes. Fixed cells were centrifuged again at 5000 g for 3 minutes, the supernatant was discarded, and the pellet was resuspended in ice-cold PBS. This washing step was repeated twice to ensure the removal of excess ethanol. Fixed cells were then processed directly for intracellular amplification or other downstream applications.

## Additional Methods Considerations

### Yeast cell culture

S. cerevisiae C5W4 WTC-SOD1-D102S was used. Cells were streaked on YP plus 2% dextrose agar plates from frozen -80°C glycerol stocks and grown at 30°C for 48 hours. Single colonies of (yeast strain) were inoculated into YP plus 2% dextrose liquid media (YPD) and grown at 30°C with shaking [200 rotations per minute (rpm)] for 24 hours.

### Bacterial cell culture

E. coli DH5α was used. Cells were streaked on LB agar plates from frozen -80°C glycerol stocks and grown at 37°C for 24 hours. Single colonies of DH5α were inoculated into fresh LB medium and grown at 37°C with shaking [200 rotations per minute (rpm)] for 24 hours.

### Cell Lysis

After amplification, the entire reaction volume was transferred into a microcentrifuge tube and 1 mL of cold PBS and 5 µL of 10% Triton X-100 was added. Samples were spun down at 4°C, 5000 g for 3 min. The supernatant was carefully aspirated off, leaving ∼ 30 µL to avoid removing the pellet. Cells were then resuspended in 1 mL cold PBS, spun down, and resuspended in 1 mL cold PBS for a total of two washes. After washing cells, the supernatant was aspirated off and cells were resuspended in 50 µL of cold PBS, 50 µL of 2X lysis buffer [20 mM Tris (pH 8.0), 400 mM NaCl, 100 mM EDTA (pH 8.0), and 4.4% SDS], and 10 µL of proteinase K solution (20 mg/mL). Cells were incubated at 55°C for 2 hours with periodic vortexing to lyse the cells and reverse the formaldehyde cross-links.

### Preparing Streptavidin Beads for Sample Binding

To prepare beads for sample binding, they must first be washed. For each lysate, 44 μL of Dynabeads MyOne Streptavidin C1 (Invitrogen) were washed three times with 800 μL of a 1X wash solution of 5 mM Tris-HCl pH 8.0, 1 M NaCl, 500 μM EDTA, and 0.05% Tween-20 using a magnetic 1.5 mL tube rack. The beads were then resuspended in 100 μL per sample of a 2X wash solution containing 10 mM Tris-HCl pH 8.0, 2 M NaCl, and 1 mM EDTA.

### Sample Binding to Streptavidin Beads

Directly after lysates were removed from heat, 5 μL of 100 μM PMSF (resuspended in isopropanol) was added to each tube and incubated at room temperature for 10 minutes to inactivate the proteinase K. To bind DNA to C1 beads, 100 μL of resuspended C1 beads were added to each sample tube and agitated at room temperature for 60 minutes. The samples were then placed on the magnetic rack and the supernatant was removed. Sampled were removed from the magnetic rack and resuspended in 250 μL of the 1X wash solution and agitated at room temperature for 5 min. Samples were replaced on the magnetic rack and the subsequent steps were repeated for a total of two wash steps. After the two 1X wash steps, the supernatant was removed, and the samples were resuspended in 250 μL of 10 mM Tris-HCl pH 8.0 and 0.1% Tween-20. At this point, the beads could be rinsed with 250 uL of molecular-grade water while the beads were still bound to the magnetic rack and moved on to subsequent steps or resuspended in 250 μL of the Tris-HCL Tween-20 buffer and stored at 4°C overnight.

### Off-the-beads Amplification

If the samples were stored overnight in the Tris-Tween buffer, the tubes were placed in a magnetic rack and the beads were rinsed with 250 μL of molecular-grade water. The bead-bound DNA was then amplified in 220 μL reactions with 2X high fidelity polymerase (KAPA HiFi) and 0.4 μM forward and reverse primers for 3 min at 95°C, and then five cycles of 98°C for 20 s, 65°C for 45 s, and 72 for 3 min.

### Quantitative PCR

The off-the-beads PCR product was then placed against a magnetic rack and the supernatant was transferred to new optical-grade PCR tubes with qPCR dye (EvaGreen 20X). The samples were then amplified on a qPCR machine for a further 10–20 cycles until the amplification curves exited the log-linear phase.

### Size selection bead clean up

The PCR products were then cleaned using a 0.8X SPRI size selection and eluted in 20 μL of molecular-grade water.

### Gel electrophoresis

5 μL of the product which was eluted during the bead clean-up was then run on a 2% agarose gel at 120 V for 15-20 min. A single properly amplified gene should appear as a dark band on the gel at the correct size of the gene. The genome that has been properly amplified with random hexamers should appear as a smear starting at approximately 5–7 kB and ending at approximately 300 bp on a gel.

## Supplemental Figures

**Supplemental Figure 1.**
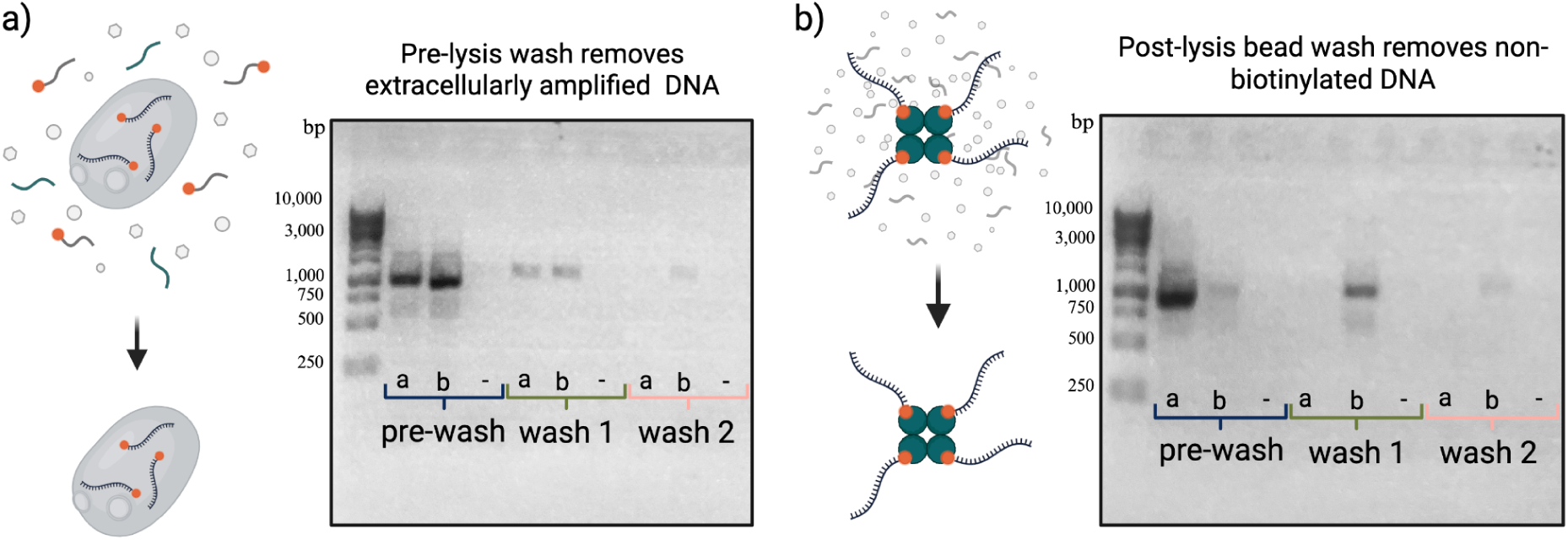
Wash steps remove extracellular and non-biotinylated DNA during the intracellular DNA amplification workflow. (a) To test whether extracellular DNA is effectively removed before lysis, biotinylated *E. coli* 16S amplicons (1 ng/μL [lane a] or 10 ng/μL [lane b]) were spiked into S. cerevisiae samples *pre-*Phi29 amplification. This is an enormous amount of extracellular DNA; 10 ng corresponds to the genomic equivalent of nearly all the yeast cells in the sample (1 million cells). This spiked-in biotinylated DNA is intended to mimic any extracellular DNA that would have been amplified with biotinylated primers by phi29. After the 16-hour incubation step during which intracellular amplification occurs, cells were subjected to two sequential pre-lysis washes. Supernatants before each wash were cleaned up using size selection beads and underwent PCR-amplification using custom *E. coli* 16S-specific primers designed to generate a ∼1 kb amplicon. The gel reveals a decreasing signal across washes, which confirms that extracellular DNA is removed by pre-lysis washes. Lane “–” is a negative control reaction with no DNA added. (b) To test whether a cell’s native genome is effectively removed after lysis and bead binding, non-biotinylated (mimicking a cell’s native genome) *E. coli* 16S amplicons (1 ng/μL [lane a] or 10 ng/μL [lane b]) were spiked into cell lysates *post* lysis. The lysates were bound to streptavidin beads in accordance with the INgen protocol. Bead-bound DNA underwent two washes. Supernatants before each wash step were cleaned up with size selection beads and PCR-amplified as in (a). The gel reveals a reduction in signal across washes, which indicates that non-biotinylated DNA is efficiently removed from bead-bound samples. Lane “–” is a negative control with no DNA added.

**Supplemental Figure 2.**
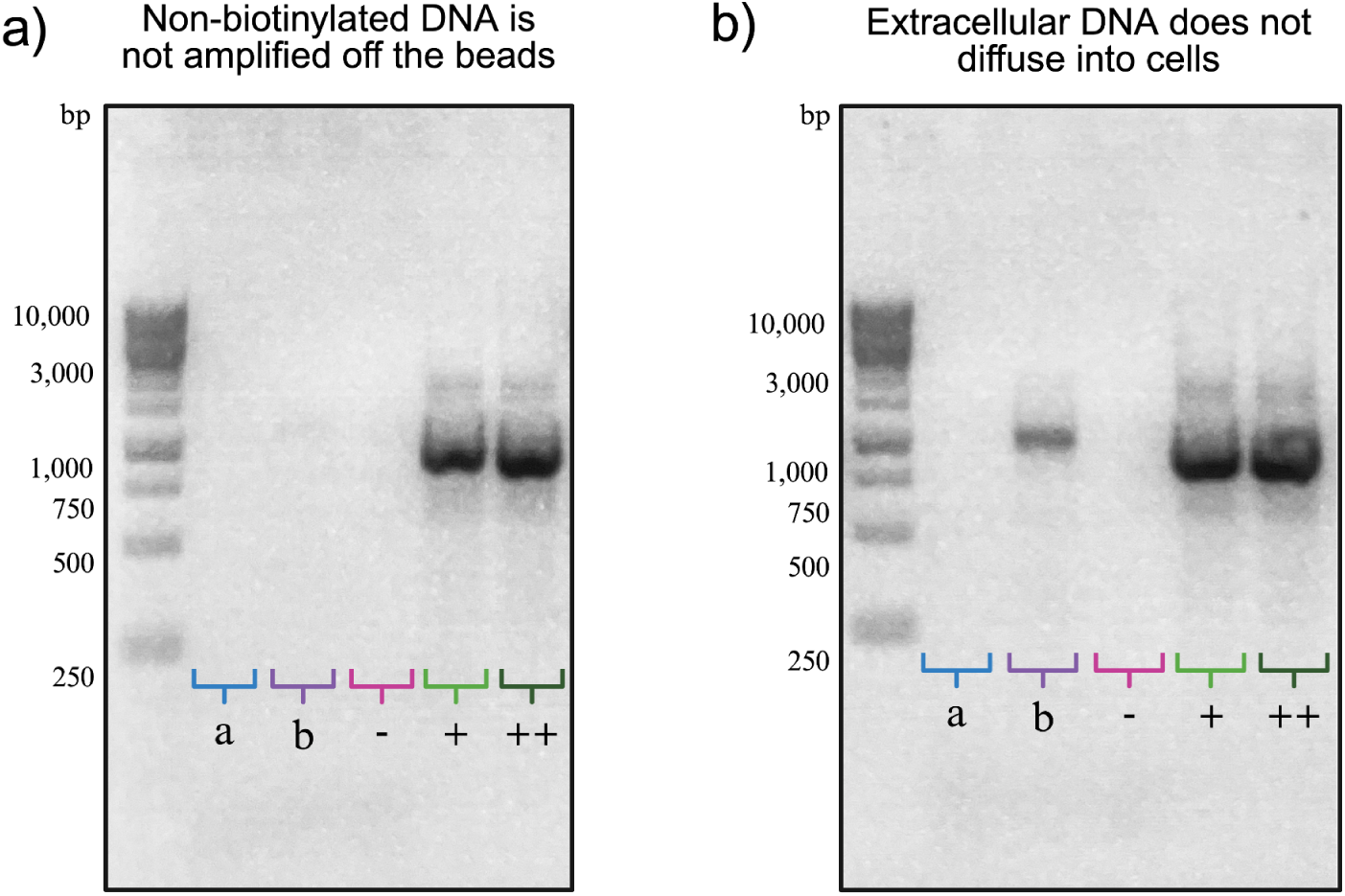
Controls demonstrating that amplification is specific to intracellular and biotinylated DNA. (a) To test whether non-biotinylated DNA binds non-specifically to beads and is retained through washing, *E. coli* 16S DNA (1 ng [lane a], 10 ng [lane b]) was spiked into lysates after cell lysis. Lysates were incubated with streptavidin beads, the beads were washed in accordance with the INgen protocol, and amplification was performed directly off the beads using *E. coli* 16S-specific primers. The absence of signal in lanes a and b demonstrates that non-biotinylated DNA does not remain associated with the beads. Lane “–” is a no-DNA negative control. Lanes “+” and “++” are positive controls where 1 ng and 10 ng of *E. coli* 16S DNA, respectively, were directly amplified using the same primers. (b) To test whether extracellular DNA can diffuse into intact cells, biotinylated *E. coli* 16S DNA (1 ng [lane a], 10 ng [lane b]) was spiked into samples before Phi29 treatment. This is an enormous amount of spiked-in DNA; 10 ng corresponds to the genomic equivalent of nearly all the yeast cells in the sample (1 million cells). After pre-lysis washes, cells were lysed and streptavidin beads were used to bind any biotinylated DNA in the lysate. Amplification off-the-beads shows no signal in lane a but some signal in lane b. These results suggest that extracellular biotinylated DNA does not appreciably diffuse into intact cells at reasonable concentrations, though it is possible with incredibly high concentrations of extracellular DNA. Lanes “–”, “+”, and “++” serve as negative and positive amplification controls as in panel (a).

**Supplemental Figure 3.**
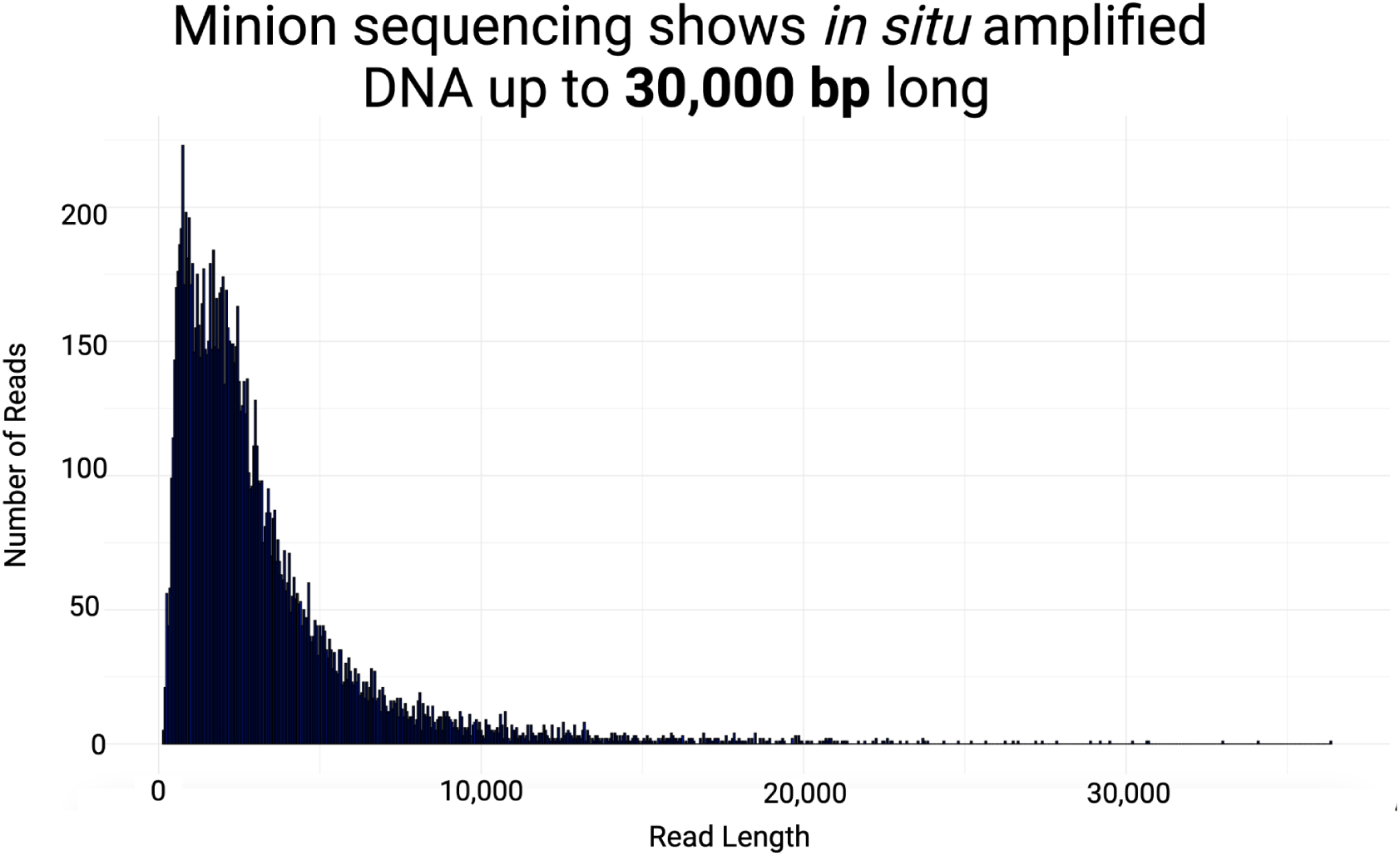
MinION sequencing shows intracellularly amplified DNA up to 30,000 bp long. Histogram of read lengths obtained from Oxford Nanopore MinION sequencing of intracellularly amplified DNA. The majority of reads are between 1,000 and 10,000 bp, with a peak around 2,000 bp. A substantial tail of longer reads extends beyond 20,000 bp, with some reads reaching lengths up to ∼30,000 bp. These data demonstrate the capability of the intracellular amplification protocol to generate long DNA products suitable for long-read sequencing.

**Supplemental Figure 4.**
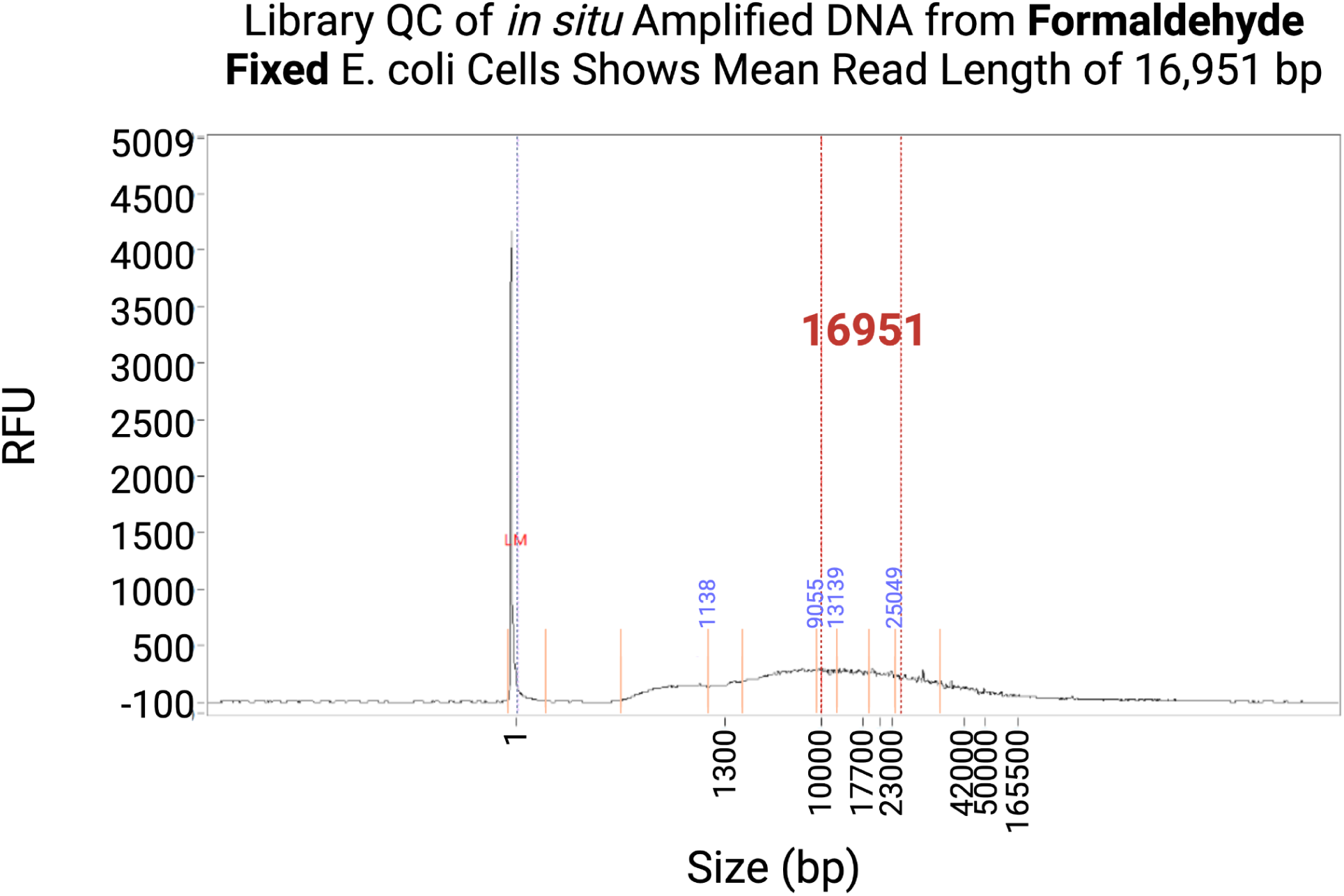
Library QC of intracellularly Amplified DNA from Formaldehyde-Fixed *E. coli* Cells Shows Mean Read Length of 16,951 bp. Femto Pulse trace of DNA amplified intracellularly from formaldehyde-fixed *E. coli* cells. The electropherogram shows a broad distribution of fragment sizes, with a mean read length of 16,951 bp, indicating successful amplification of long DNA molecules.

**Supplemental Figure 5.**
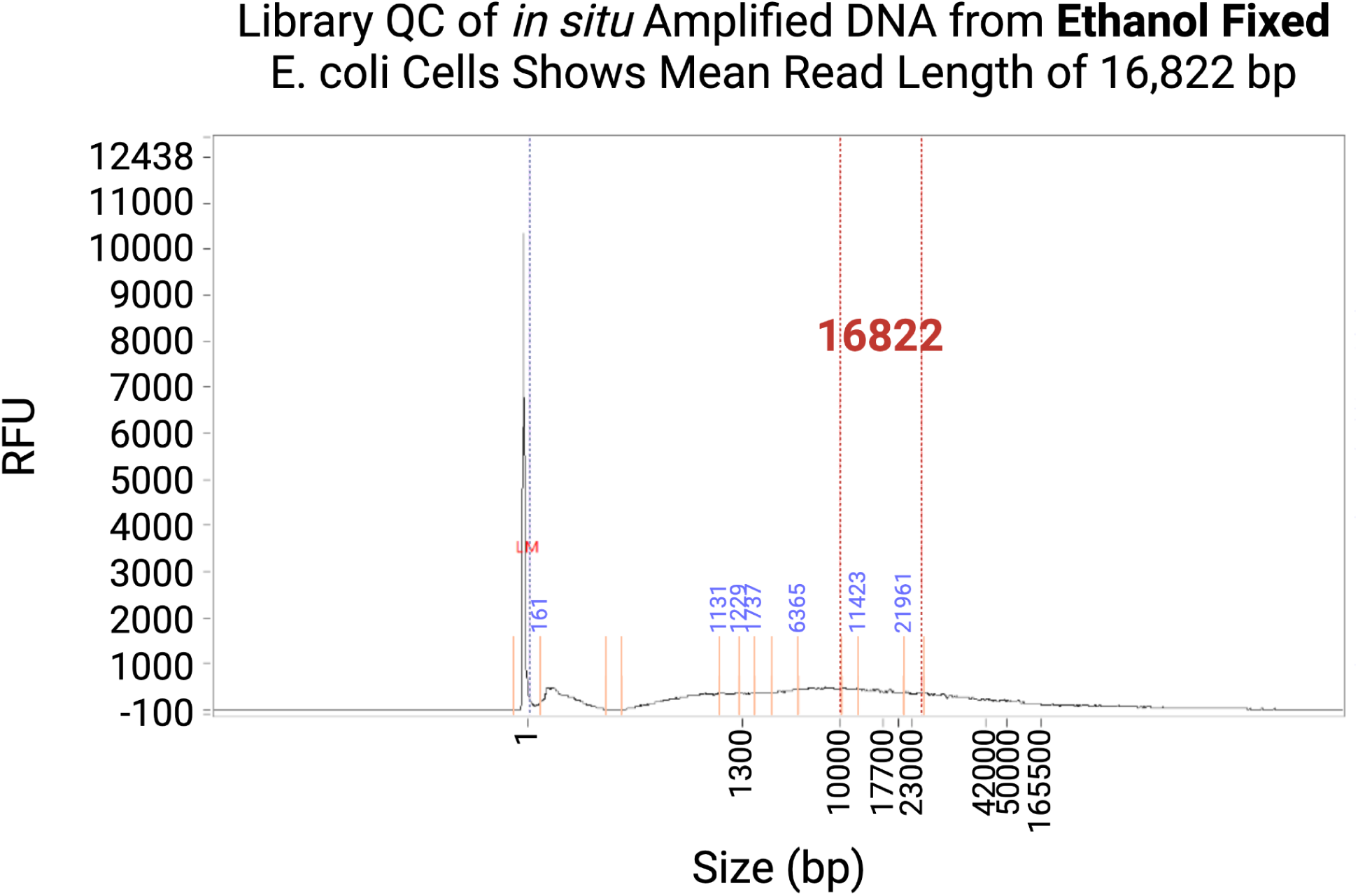
Library QC of intracellularly Amplified DNA from Ethanol-Fixed *E. coli* Cells Shows Mean Read Length of 16,822 bp. Femto Pulse trace of DNA amplified intracellularly from ethanol-fixed *E. coli* cells. The size profile closely resembles that of the formaldehyde-fixed sample, with a mean fragment length of 16,822 bp, supporting robust amplification from both fixation conditions.

**Supplemental Figure 6.**
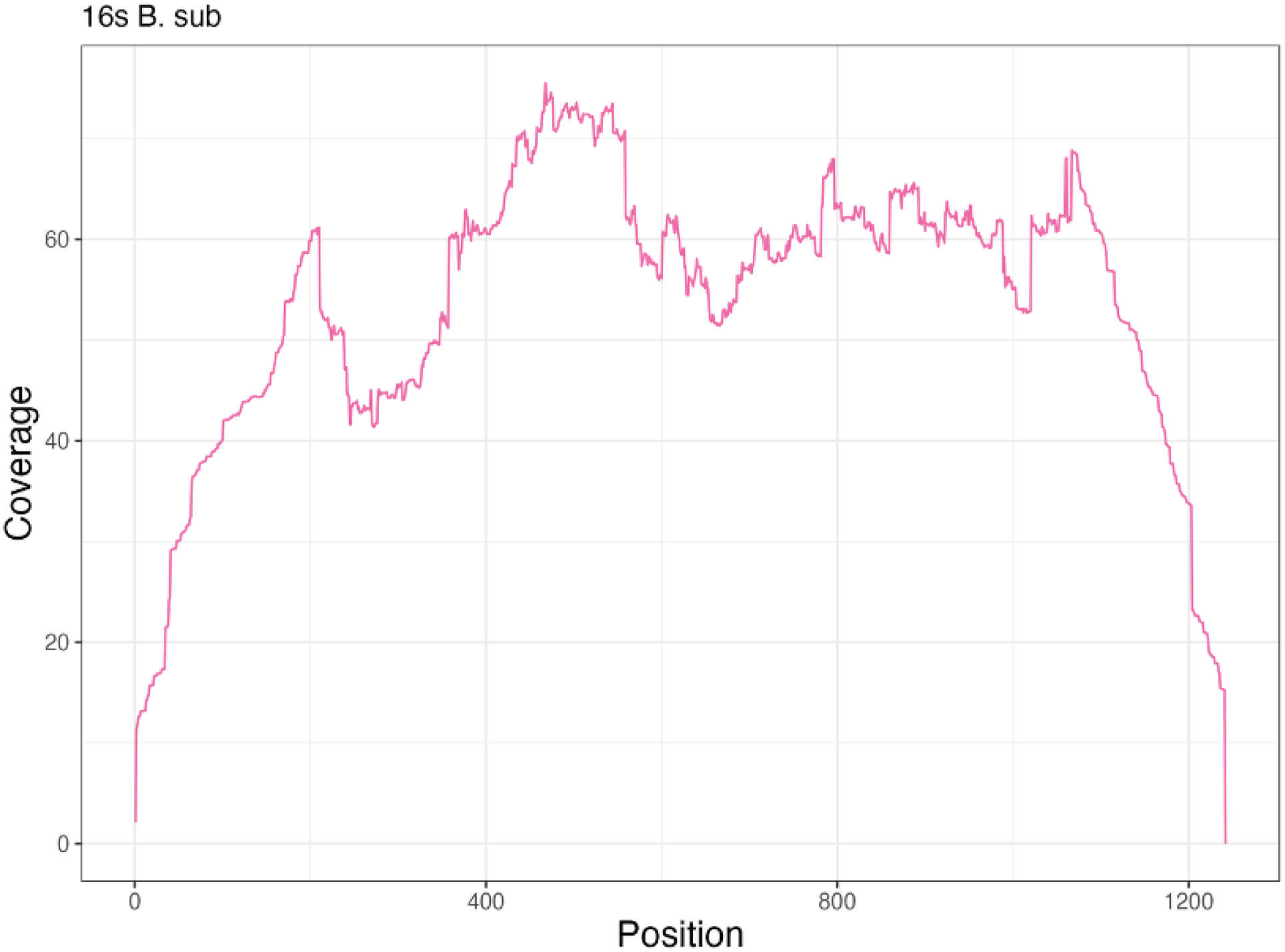
Coverage across the 16S rRNA gene of *Bacillus subtilis* following intracellular amplification and sequencing. Coverage plot showing read depth across the length of the *B. subtilis* 16S rRNA gene. DNA was first amplified intracellularly using gene-specific primers and Phi29 polymerase, followed by lysis and streptavidin bead capture. The bead-bound product was then re-amplified using specific primers and OneTaq polymerase. This off-the-beads product was sequenced, and the resulting reads demonstrate high coverage across the gene, with expected tapering at the 5’ and 3’ ends. These results indicate efficient and relatively uniform amplification of full-length 16S rRNA sequences using this workflow.

